## Supplemental Figures for "Potent pollen gene regulation by DNA glycosylases in maize"

### Supplementary Figures

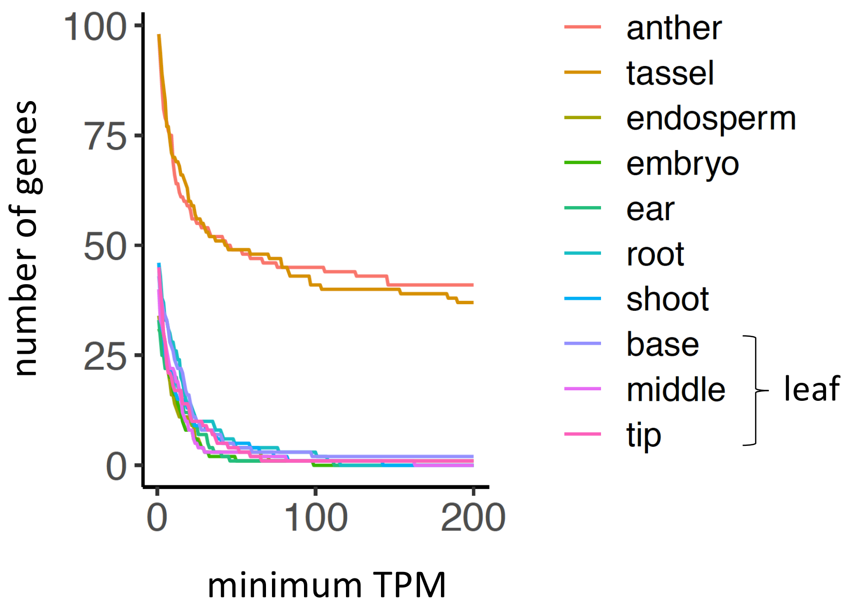


**Supplemental Figure 1**

The Y axis indicates the number of genes with TE-like methylation, plotted relative to the minimum expression thresholds (X axis) in each tissue type. This analysis only includes genes that are present at syntenic positions in B73 and the other 25 NAM founder genomes (core genes) and whose coding DNA sequence (CDS) does not overlap at all with TE annotations.

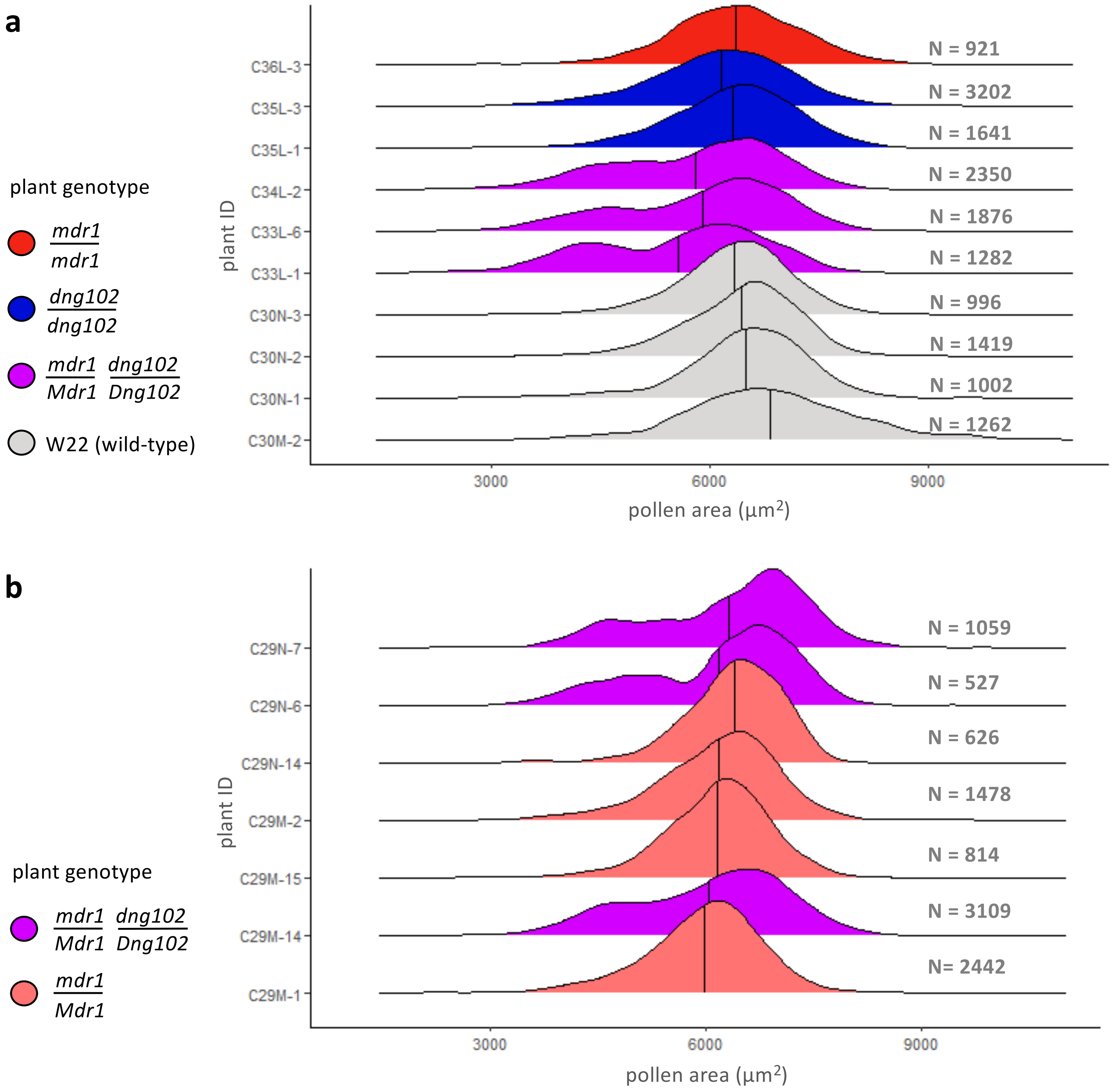


**Supplemental Figure 2**

**(A)** Distribution of 2-D pollen grain areas based on microscope image analysis. Each distribution is derived from pollen grains from a single plant whose genotype is indicated by the color on the left. All plants are near isogenic with W22. Plant IDs are in the format "family"-"sibling number". Capital allele indicates WT, lowercase mutant. The *mdr1* allele is the original Kermicle allele, and *dng102* allele is the Q235 allele. The number of pollen grains analyzed for each plant is indicated to the right.

**(B)** Distribution of 2-D pollen grain areas based on microscope image analysis from plants that are near isogenic with W22, as in A. All plants are siblings. The *mdr1* allele is EMS4-06835d, and *dng102* is the Q235 allele.

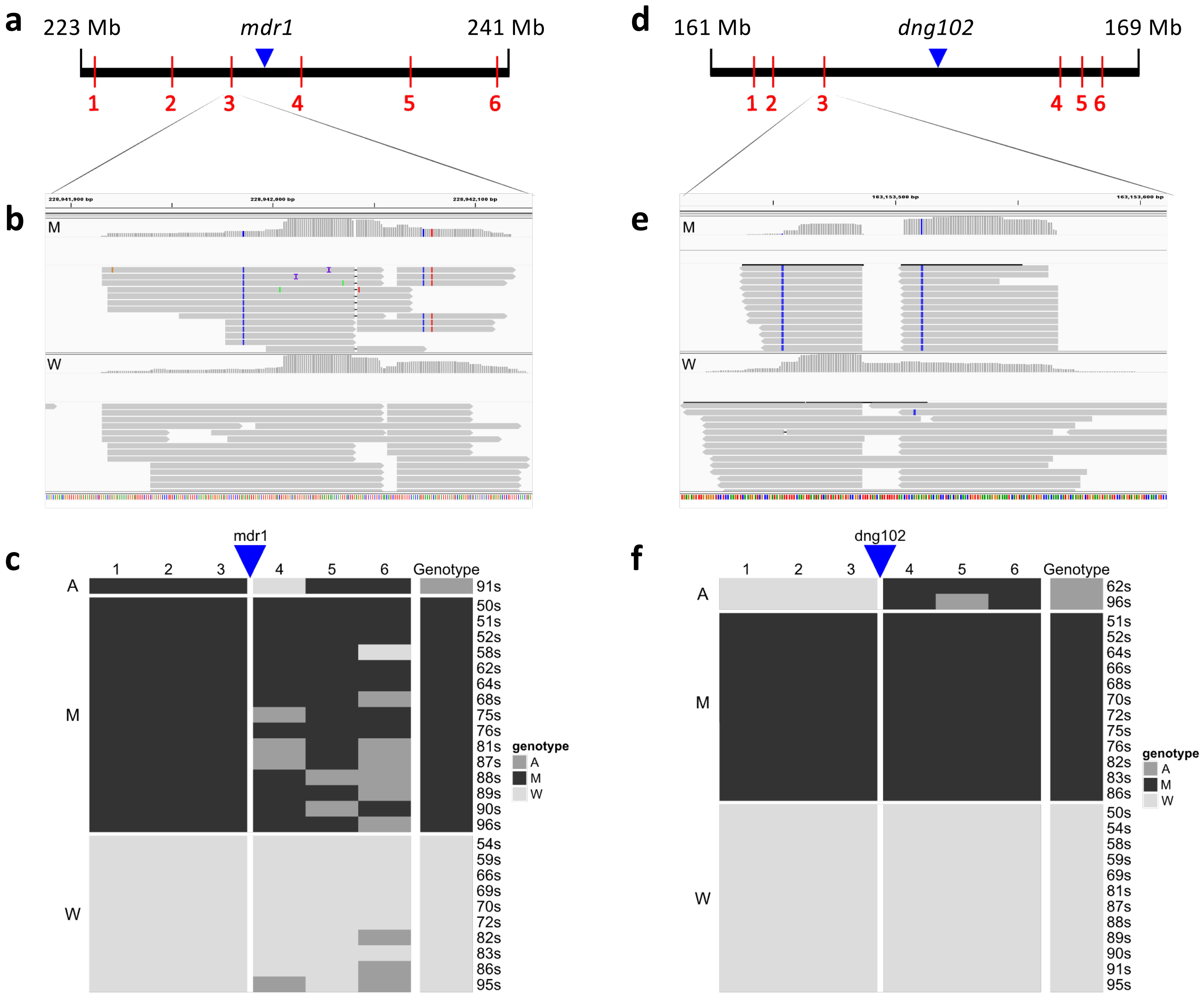


### Supplemental Figure 3

**(A, D)** Map of DNG and sentinel gene locations on chromosome 4 (*mdr1*) (A) and chromosome 5 (*dng102*) (D). *mdr1*-linked sentinel genes are (1) Zm00004b023798, (2) Zm00004b023832, (3) Zm00004b023880, (4) Zm00004b023919, (5) Zm00004b024000, and (6) Zm00004b024077. *dng102*-linked sentinel genes are (1) Zm00004b013764, (2) Zm00004b013769, (3) Zm00004b013785, (4) Zm00004b013866, (5) Zm00004b013875, and (6) Zm00004b013879. Sentinel genes were inherited from B73 along with *mdr1* and *dng102* and have sufficient expression in pollen to detect SNPs relative to the W22 reference genome. WT *Mdr1* and *Dng102* alleles and the rest of the genome are from W22.

**(B, E)** IGV browser images showing B73 SNPs (marked blue and red) in sequencing reads (gray arrows) at example sentinel genes in mutant (M) but not in WT (W), linked to *mdr1* (B) or *dng102* (E).

**(C, F)** Summary of pollen grain genotypes at each DNG locus, (C) for *mdr1*, (F) for *dng102*, based on SNP calling (left, M = mutant; W = wild-type). Each row represents a single pollen transcriptome, each column a sentinel gene. Genotypes at sentinel genes: black is mutant (M), light grey is wild-type (W). In cases where no reads were detected at the SNP sites at a sentinel gene, it was disregarded for that transcriptome (dark gray, A = ambiguous). In three pollen grains, the sentinel genes nearest to *dng102* or *mdr1* had conflicting genotypes, possibly due to recombination between the sentinel genes; these pollen grains were scored as ambiguous genotypes at the respective loci.

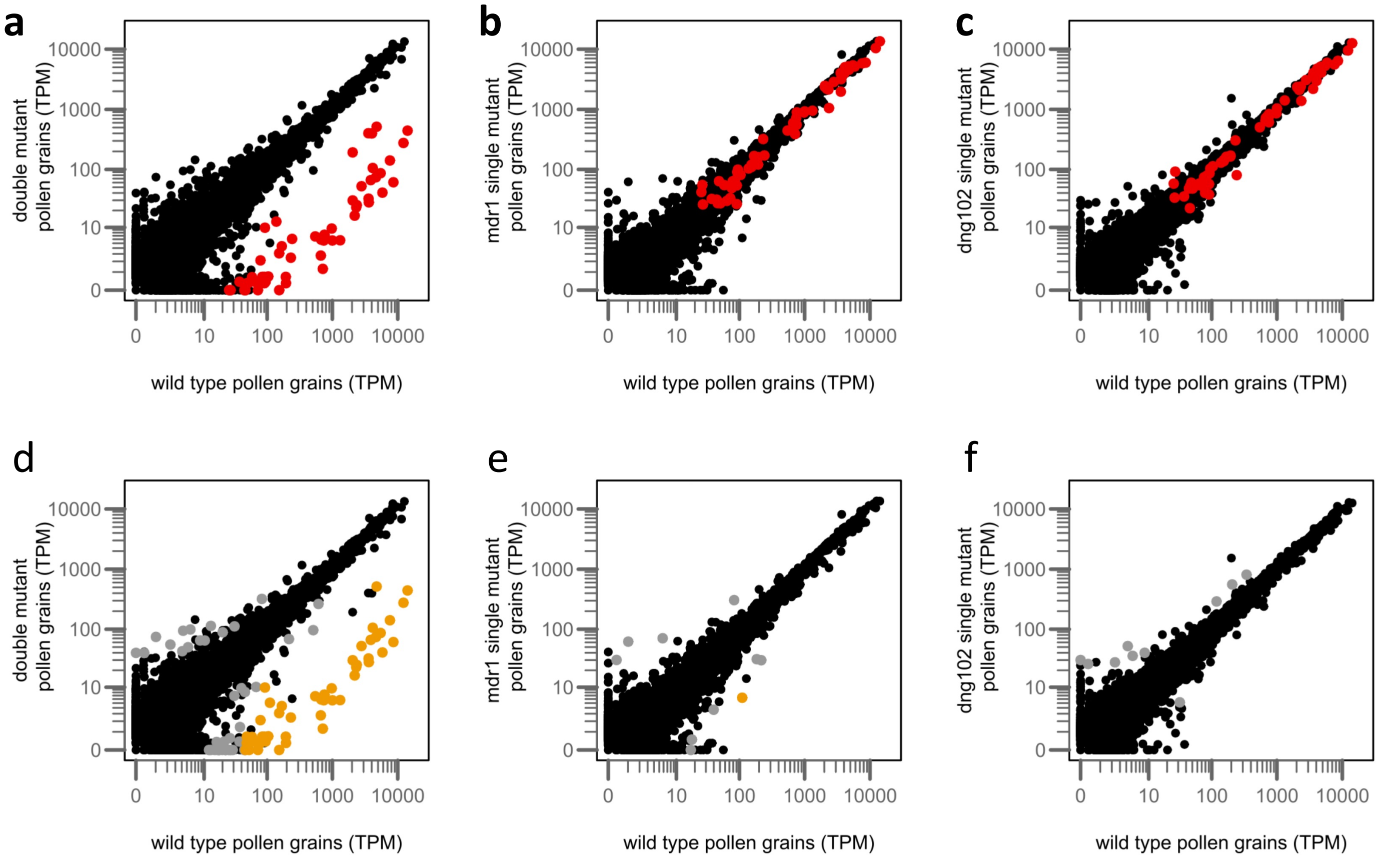


### Supplemental Figure 4

**(A-C).** Expression of DEGs (highlighted in red) from Figure 2A, plotted in WT vs. double mutant (A); WT vs. *mdr1* single mutant (B); and WT vs. *dng102* single mutant (C).

**(D-F)**. Differential expression of individual genes in WT vs. double mutant (D), WT vs. *mdr1* single mutant (E), and WT vs. *dng102* single mutant (F). Strongly differentially expressed genes (adjusted p-value ≤ 0.05; ≥ 8-fold change in expression; ≥ 10 average UMI counts, as estimated by DESeq2) are highlighted in orange. Weakly differentially expressed genes (adjusted p-value ≤ 0.05; ≥ 2-fold change in expression) are highlighted in gray.

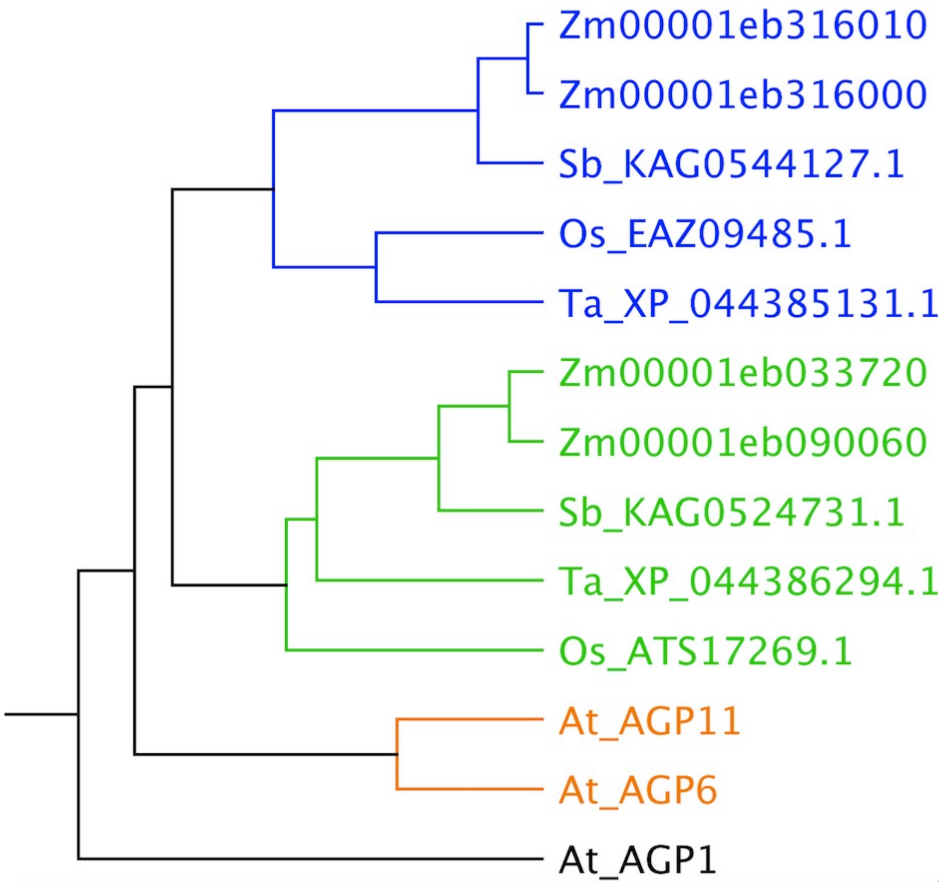


**Supplemental Figure 5**

Protein tree of arabinogalactan protein (AGP) homologs, generated using UPGMA based on sequence similarity. Zm indicates *Zea mays*, Sb *Sorghum bicolor*, Os *Oryza sativa*, Ta *Triticum aestivum,* and At *Arabidopsis thaliana.*

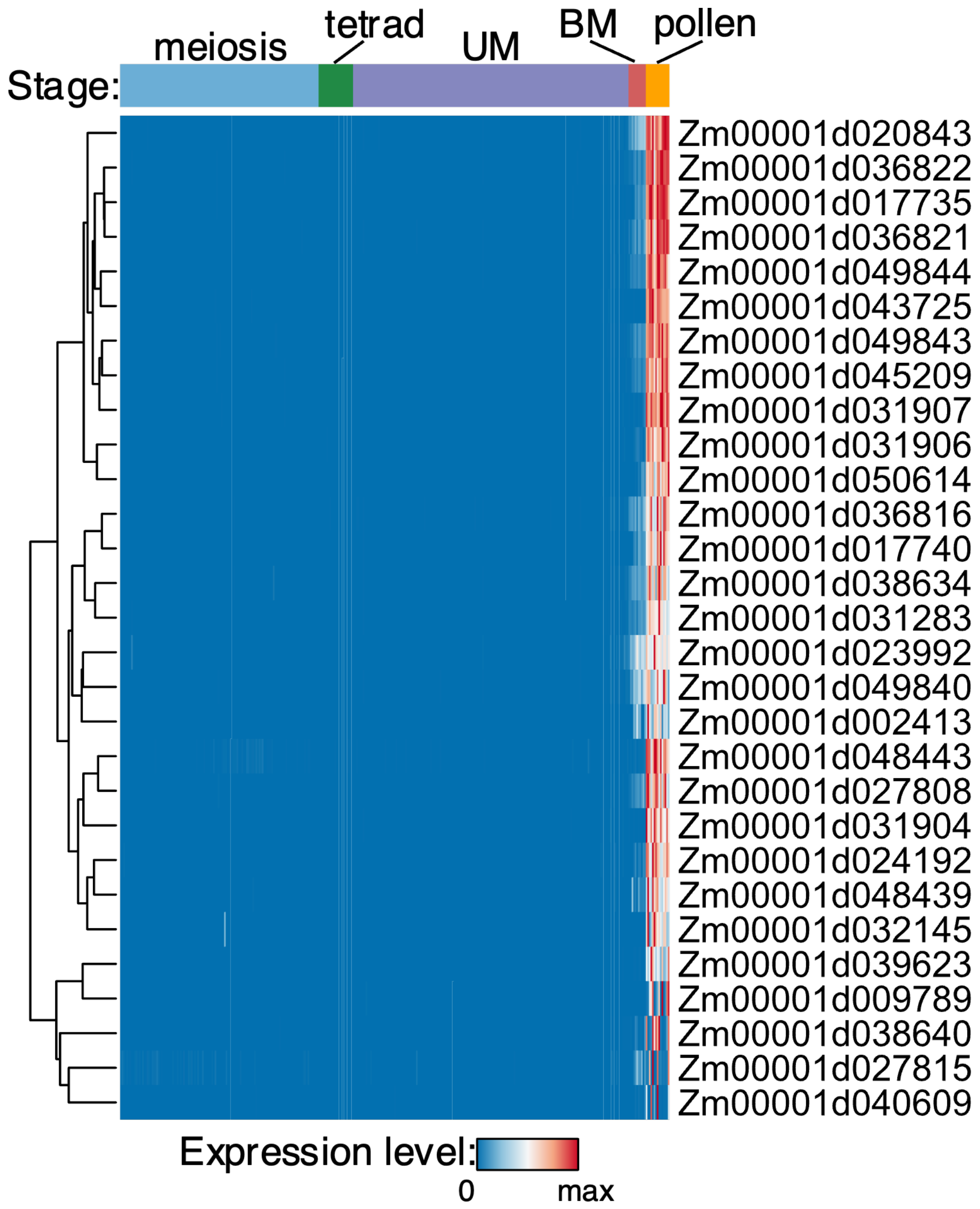


**Supplemental Figure 6**

Expression timecourse of DEGs. UM = unicellular microspore, BM = bicellular microspore. Data are from B73 single-cell transcriptomes (Nelms and Walbot 2022). Each column represents a single pollen grain or pollen precursor. Gene names are listed as B73 v4 names because the published expression analysis was carried with v4 annotations. Mappings used between v4 and v5 are shown in Supplemental Table 3.

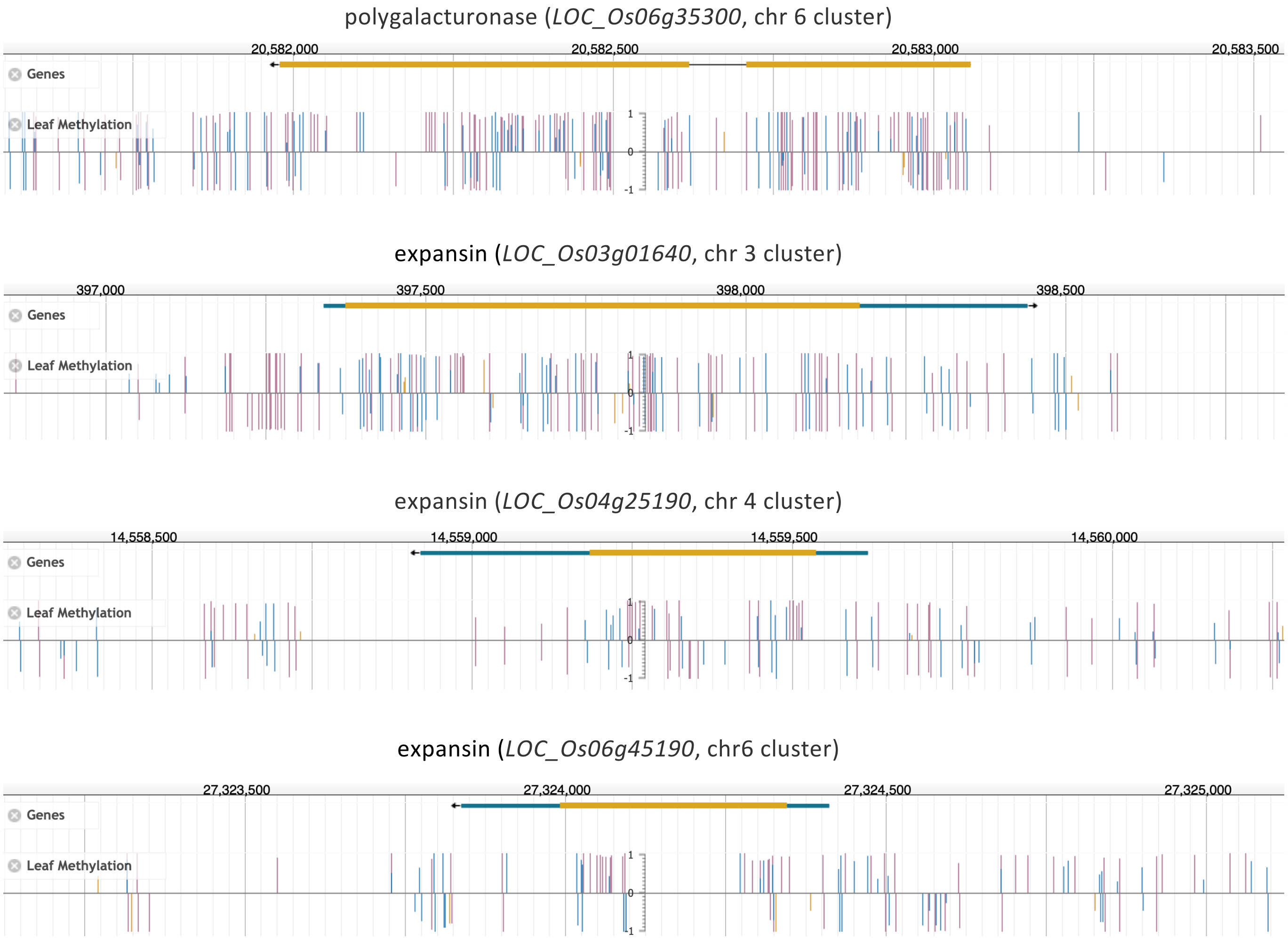


**Supplemental Figure 7**

Plant Epigenome Browser 46 showing DNA methylation over 2 Kb regions including previously identified clustered pollen genes. CG methylation is indicated in magenta, CHG in blue, and CHH in orange. Reference coordinates are based on the Os-Nipponbare-Reference-IRGSP-1.0 assembly, v7 annotations.

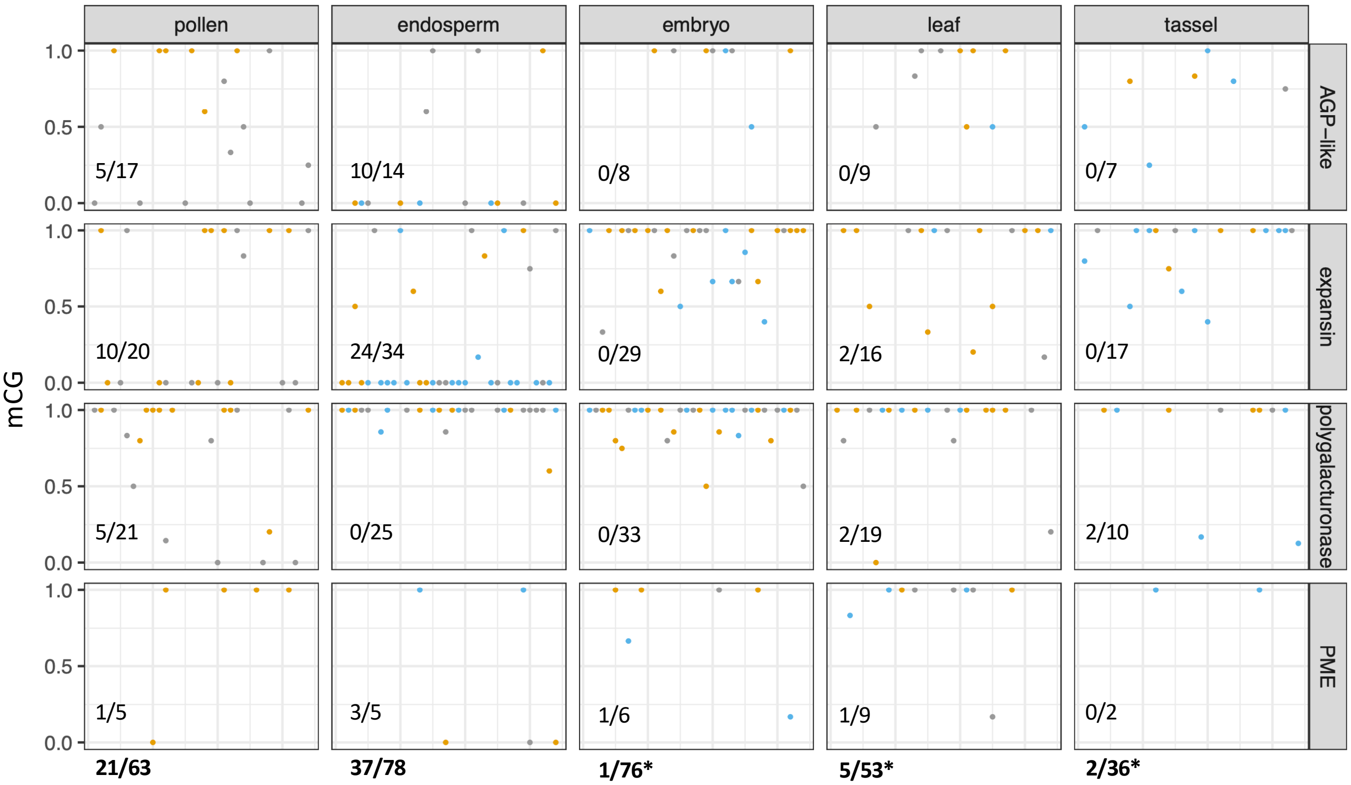


**Supplemental Figure 8**
Single-read CG methylation calls from four MPG promoters. Each dot represents a read or read segment, biological replicates distinguished by color. All reads or segments of reads that overlapped 600 bp upstream of each gene in the W22 genome and which included at least four CGs were included in the analysis. The AGP-like gene corresponds to Zm00001eb316000, the expansin to Zm00001eb130160, the polygalacturonase to Zm00001eb132550, and the pectin methylesterase (PME) to Zm00001eb239290. Numbers in the bottom left of each panel indicate proportion of reads with less than or equal to CG methylation values of 0.2.
